# SNP-level *F_ST_* outperforms window statistics for detecting soft sweeps in local adaptation

**DOI:** 10.1101/2022.01.12.476036

**Authors:** Tiago da Silva Ribeiro, José A. Galván, John E. Pool

**Affiliations:** Department of Integrative Biology, University of Wisconsin-Madison, Madison, WI, 53706, USA; Laboratory of Genetics, University of Wisconsin-Madison, Madison, WI, 53706, USA; John Jay College of Criminal Justice, New York, NY, 10019, USA

**Keywords:** Local adaptation, soft sweeps, partial sweeps, population genomics, *Drosophila melanogaster*

## Abstract

Local adaptation can lead to elevated genetic differentiation at the targeted genetic variant and nearby sites. Selective sweeps come in different forms, and depending on the initial and final frequencies of a favored variant, very different patterns of genetic variation may be produced. If local selection favors an existing variant that had already recombined onto multiple genetic backgrounds, then the width of elevated genetic differentiation (high *F_ST_*) may be too narrow to detect using a typical windowed genome scan, even if the targeted variant becomes highly differentiated. We therefore used a simulation approach to investigate the power of SNP-level *F_ST_* (specifically, the maximum SNP *F_ST_* value within a window) to detect diverse scenarios of local adaptation, and compared it against whole-window *F_ST_* and the Comparative Haplotype Identity statistic. We found that SNP *F_ST_* had superior power to detect complete or mostly complete soft sweeps, but lesser power than window-wide statistics to detect partial hard sweeps. To investigate the relative enrichment and nature of SNP *F_ST_* outliers from real data, we applied the two *F_ST_* statistics to a panel of *Drosophila melanogaster* populations. We found that SNP *F_ST_* had a genome-wide enrichment of outliers compared to demographic expectations, and though it yielded a lesser enrichment than window *F_ST_*, it detected mostly unique outlier genes and functional categories. Our results suggest that SNP *F_ST_* is highly complementary to typical window-based approaches for detecting local adaptation, and merits inclusion in future genome scans and methodologies.

**Significance statement:** Studies that use genetic variation to search for genes evolving under population-specific natural selection tend to analyze data at the level of genomic windows that may each contain hundreds of variable sites. However, some models of natural selection (*e.g.* favoring an existing genetic variant) may result in genetic signals of local adaptation that are too narrow to be detected by such approaches. Here we use both simulations and empirical data analysis to show that searching for a site-specific signal of elevated genetic differentiation can find instances of local adaptation that other approaches miss, and therefore the integration of this signal into future studies may significantly improve our understanding of adaptive evolution and its genetic targets.

## Introduction

Geographically distinct populations are exposed to different selective pressures, which may result in local adaptation. The detection of genomic regions under positive selection specific to one population is essential to uncovering the genetic basis of locally adaptive trait variation.

Local adaptation can exist between populations with low genome-wide genetic differentiation, and comparing genetic variation between these closely-related populations can allow for much more powerful detection of positive selection than is possible from a single population. In light of that advantage, as well as the potential applicability of genetic mapping and functional approaches to locally adaptive traits, local adaptation has played a key role in our increasing understanding of adaptive evolution at the genetic level (Kawecki and Ebert 2004; Yeaman 2015; Tigano and Friesen 2016). In addition to its importance for evolutionary biology and ecology, the identification of regions under selection has implications for applied fields such as health sciences and agriculture because it can also pinpoint regions of the genome that hold functional diversity (Bamshad and Wooding 2003; Ross-Ibarra *et al*. 2007). There has also been increasing recognition of the importance of local adaptation for a species’ future adaptive potential, with implications for conservation genetics and adaptation to climate change (Funk *et al*. 2012; Aitken and Whitlock 2013; Fitzpatrick and Keller 2015).

Population genomic scans for local adaptation compare genetic variation between two populations, often searching for specific genomic windows that depart from genome-wide patterns of differentiation in a manner consistent with population-specific natural selection. Positive selection has traditionally been conceptualized and modeled as a selective sweep, which traditionally involves a new beneficial mutation rising to fixation, with strong effects on genetic variation at linked sites (Maynard Smith and Haigh 1974; Kaplan *et al*. 1989). However, there are different kinds of selective sweeps, depending on the initial and final frequencies of the favored variant, and different statistical tests for deviations from neutrality vary in their power to detect them.

First, selective sweeps can be classified as hard or soft sweeps. In a hard sweep, only a single original haplotype carrying the advantageous allele is boosted by natural selection. This situation might be expected if selection favors either a newly occurring mutation or else a variant at low enough frequency that only one copy contributes to the sweep by chance. In a soft sweep, two or more distinct haplotypes carrying the beneficial variant increase in frequency. In some cases, soft sweeps occur because the advantageous allele was present in the population, segregating neutrally, prior to the onset of selection (Hermisson and Pennings 2005). But they can also be the result of recurrent mutations or influx of new alleles through migration (Pennings and Hermisson 2006a, 2006b).

Selective sweeps can also be classified as complete or partial sweeps. In a complete sweep, the advantageous allele reaches fixation in the population. In a partial sweep, the advantageous allele is at an intermediary frequency. This may occur either because the sweep is still ongoing or because positive selection ended prior to fixation. Situations in which a sweep might terminate prematurely include an environmental change, a polygenic trait reaching its new optimum or threshold value, or an allele reaching a balanced equilibrium in a scenario such as heterozygote advantage.

Different kinds of selective sweeps leave different signatures of local adaptation and our power to detect them will differ depending on which methods we use (Lange and Pool 2016).Some common approaches to scanning the genome for population-specific selective sweeps use *F_ST_* (or *F_ST_*-based) statistics to quantify genetic differentiation between populations. Local adaptation is expected to create genomic regions with higher differentiation than what would be expected under neutrality, since allele frequencies in these regions will change faster as the beneficial allele increases in frequency (Lewontin and Krakauer 1973). Neutral expectations can be inferred either with demographic simulations or an outlier approach. Demographic simulations, based on a previously estimated model of population history, can be used to mimic the history of the populations being studied in the absence of natural selection. Outlier approaches rely on the genome-wide distribution of *F_ST_* as a proxy for the neutral distribution, since neutral forces (including those due to demographic history) can broadly be expected to affect the whole genome similarly. Genome scans for regions under selection have typically focused on measuring *F_ST_* or other statistics in windows of the genome of some predefined size to search for highly differentiated genomic regions.

A motivating empirical example for the present study comes from an investigation of the genetic basis of locally adaptive melanism in high altitude *Drosophila melanogaster* populations. Here, the authors used QTL mapping to identify genomic regions associated with derived dark pigmentation traits, and then used *F_ST_* to scan these regions for signatures of selection (Bastide *et al*. 2016). One very narrow and strong QTL for highland Ethiopian melanism contained the well-known pigmentation gene *ebony*, which also contributed to melanic evolution in a Uganda population (Pool and Aquadro 2007; Rebeiz *et al*. 2009). Assessing genetic differentiation between the Ethiopia and Zambia populations for the window containing *ebony*, although window-wide *F_ST_* was only marginally elevated, it had a SNP with extremely high *F_ST_* (0.85). Compared to demographic simulations, this window’s maximum SNP *F_ST_* value was among the top 1% of all windows, while its window-wide *F_ST_* was only among the 7% highest (Bastide *et al*. 2016). Simulated scenarios of soft sweeps from standing variation replicated this pattern of extremely high SNP *F_ST_* and only moderately high window *F_ST_*, suggesting that some kinds of selective sweeps that may not be detected using window-wide *F*_ST_ could potentially be detected with a SNP-level *F_ST_* approach. Further potential support for the use of SNP *F*_ST_ to detect adaptive events in this same species is demonstrated by much stronger parallel signatures of selection seen at the SNP level compared to the window level in populations that independently adapted to cold environments (Pool *et al*. 2017).

Challenges of using SNP *F_ST_* values include their variability due to random sampling effects (Weir *et al*. 2005) and the large number of tests that need to be made against a null distribution. Therefore, larger sample sizes are needed than for window *F_ST_*. By using only the highest SNP *F_ST_* value within a window, and comparing against null simulations with demography and recombination, we may somewhat improve the multiple testing issue, since here we are not treating all tightly linked SNPs as fully independent tests. Another advantage of this approach is to make a SNP *F_ST_* genome scan easier to compare to window-wide statistics. If these metrics are able to detect different types of selective events, then a more comprehensive scan for signatures of selection could benefit from using both window and SNP-based methods. The genome-wide distribution of these statistics in natural populations, compared to their neutral expectations, might also shed light on the contribution of different kinds of selective sweeps to local adaptation.

To understand the utility of using the highest *F_ST_* value of any SNP within a window (herein *F_ST_MaxSNP_*) as a local adaptation summary statistic, we performed power analyses based on extensive simulations, and then applied these results to empirical data from natural populations of *D. melanogaster*. We calculated the power of *F_ST_MaxSNP_* to detect signatures of local adaptation under a wide range of different selective scenarios and demographic histories. We performed demographic simulations and compared the power of *F_ST_MaxSNP_* to both window-wide *F_ST_* (herein, *F_ST_Window_*) and a comparative haplotype-based statistic (*χ_MD_*). Then, we investigated the genome-wide distribution of *F_ST_MaxSNP_* and *F_ST_Window_* among several natural populations of *D. melanogaster*, to determine whether either statistic was enriched genome-wide in empirical data compared to neutral expectations. Finally, we used an outlier approach to perform a genome scan for regions potentially under local adaptation between the Ethiopia and Zambia populations mentioned above, using *F_ST_MaxSNP_*, *F_ST_Window_*, and *χ_MD_*, and we determined the extent of overlap between candidate regions identified according to these different methods. These analyses allowed us to both identify the parameter space in which *F_ST_MaxSNP_* outperforms other statistics, and to assess the utility and complementarity of applying these approaches to real data.

## Results

### SNP-level *F_ST_* and window-wide summaries have complementary power to detect local adaptation

We performed power analyses of *F_ST_MaxSNP_*, *F_ST_Window_*, and *χ_MD_* using population genetic simulations with and without natural selection. We used *msms* (Ewing and Hermisson 2010) to simulate population-specific selective sweeps with constrained initial and final allele frequencies, as well as scenarios with population size bottlenecks or migration (simulation commands in Table S1). For each scenario, we simulated populations with high effective population size (*N_e_*) using a set of parameters based on *D. melanogaster* and populations with low *N_e_* using parameters based on humans, following the design of a previous power analysis study that did not include *F_ST_MaxSNP_* (Lange and Pool 2016). Power was defined in a locus-specific context, based on the proportion of selection simulations giving a more extreme value of the summary statistic than the 95th quantile of its distribution from neutral simulations.

Unsurprisingly, all three statistics were found to have high power for the case of complete hard sweeps (Figure 1; Table S1). These simulations were conditioned on fixation of a beneficial new mutation in one population that had not occurred in the other population. In light of this fixed difference, *F_ST_MaxSNP_* in all replicates had its maximum value (*F_ST_MaxSNP_* = 1). In such cases, the power of *F_ST_MaxSNP_* was binary, either zero or one, depending on whether or not 5% of the corresponding neutral replicates had an allele that reached fixation. In our simple isolation model, the likelihood that a neutral allele can reach fixation increases with the split time (Table S1; Figure S1). Stronger bottlenecks also boost the likelihood of having neutral alleles reach fixation (Table S1; Figure S2, Figure S3). Hence, power for *F_ST_MaxSNP_* to detect complete hard sweeps goes from high, for recent splits and weaker bottlenecks, to zero for histories in which more than 5% of neutral replicates contain a fixed difference. Similarly, *F_ST_Window_* and *χ_MD_* had higher power to detect signatures of local adaptation following recent splits and in weaker bottlenecks, but their change in power was gradual and continuous instead of binary.

**Figure 1.**
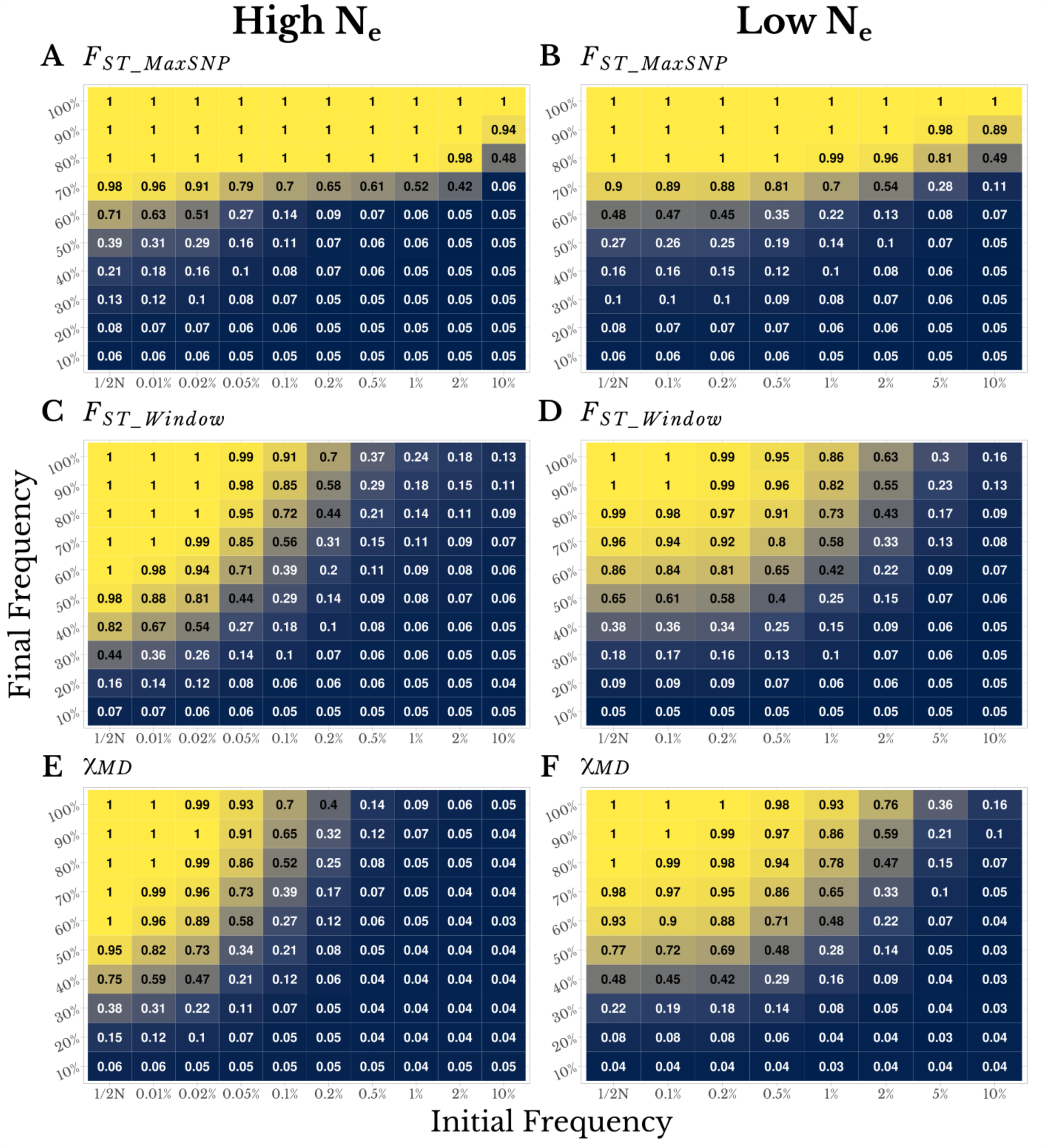
SNP-level *F_ST_* and window-wide statistics show complementary power to detect local adaptation, depending on the type of selective sweep simulated. Numbers and colors in each panel both depict statistical power to detect local adaptation, in high *N_e_* populations (s=0.001, left column) and low N_e_ populations (s=0.01, right column). In each panel, the x-axis illustrates the pre-selection frequency of a favored variant (with the left column indicating selection on newly-occurring mutations) and the y-axis illustrates the final frequency of the sweep (with the top row showing complete sweeps). Detection power is shown for (A and D) *F_ST_MaxSNP_*, (B and E) *F_ST_Window_*, and (C and F) *χ_MD_*. These results are based on a demographic history of simple isolation between two populations without change in population size, with a split time of 0.2*N_e_* generations.

In the case of complete or nearly complete soft sweeps, *F_ST_MaxSNP_* showed a clear power advantage over *F_ST_Window_* and *χ_MD_*. Notably, for sweeps ending between 80% and 100% frequency, *F_ST_MaxSNP_* had high power to detect local adaptation, even for cases with rather high initial frequencies of the beneficial allele (e.g. 10%; Figure 1; Figure 2). In contrast, *F_ST_Window_* and *χ_MD_* showed rapidly diminishing performance as sweeps became softer (Figure 1; Figure 2). These results make sense, in that beneficial alleles that drift to higher pre-selection frequencies have more time to recombine onto multiple haplotypes, and recombination events will have happened closer to the selected site on average. Therefore, soft sweeps are generally narrower in width and may not substantially alter window-wide statistics (Catania *et al*. 2004; Schlenke and Begun 2004; Hermisson and Pennings 2005). Although the two window-wide statistics maintained good power for lower initial frequencies, some of the replicates of those scenarios are actually generating hard sweeps due to the chance survival of a single haplotype carrying the favored variant (Jensen 2014), as shown by an average number of beneficial haplotypes lower than two in these simulations (Figure 2). Moreover, as the average number of haplotypes carrying the favored variant increased, the power of the window-wide statistics decreased (Figure 2), while the power of *F_ST_MaxSNP_* was unchanged.

**Figure 2.**
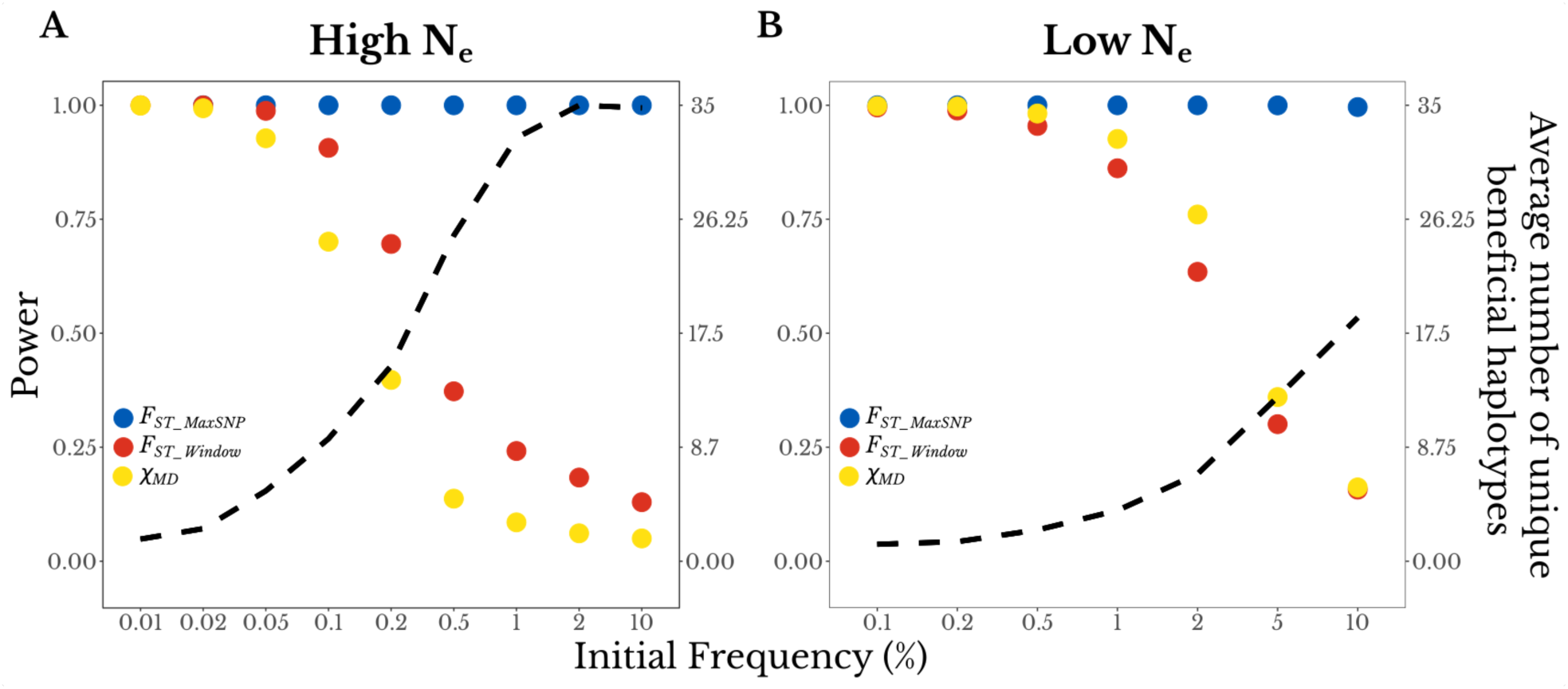
*F_ST_MaxSNP_* shows an increasing power advantage as sweeps become softer. For complete sweeps with a range of initial frequencies (x-axis), the two y-axes show detection power for each statistic (left axis, dots) and the average number of unique beneficial haplotypes present at the end of the simulation (right axis, dashed line). Results are shown for (A) high *N_e_* populations (s=0.001) and (B) low N_e_ populations (s=0.01), for the same demographic history as in Figure 1.

Contrasting results were obtained for partial, harder sweep scenarios. In cases where new mutations or rare standing variants were only boosted to intermediate frequencies, *F_ST_Window_* and *χ_MD_* had fairly strong power, whereas *F_ST_MaxSNP_* declined sharply in effectiveness at around 60% final frequency for hard sweeps (Figure 1). These results are also intuitive, in that partial hard sweeps can meaningfully alter allele frequencies across a whole window and generate a class of identical haplotypes, even though no single SNP traverses an extreme range of frequencies. The broadly similar power profiles of *F_ST_Window_* and *χ_MD_* are somewhat surprising in light of their distinct basis (albeit consistent with Lange and Pool, 2016). Less surprising is that for the challenging scenario of partial soft sweeps, none of the three statistics showed strong power in the scenarios examined (Figure 1).

Whereas the above simulations had no migration, we also wondered if *F_ST_MaxSNP_* might prove useful in detecting targets of local adaptation for which genetic differentiation had been whittled down in width by recombination with migrant alleles over time. We therefore simulated scenarios with varying combinations of migration rate and population split time, while assuming symmetric migration rates and equal but opposing selective pressures. Overall, *F_ST_MaxSNP_* and *F_ST_Window_* performed very similarly to each other and better than *χ_MD_*. Particularly in the high *N_e_* scenarios (which feature a higher ratio of recombination to mutation events) with intermediate migration rates, there was a narrow space of parameters in which *F_ST_MaxSNP_* performed slightly better than *F_ST_Window_* (Figure S4). The split time between the populations greatly affected the power of *χ_MD_*, which performed better on recent splits. The power of the *F_ST_* statistics showed a small improvement for more recent splits and intermediate migration rates. Although small, the effect of split time also seemed more pronounced on *F_ST_Window_* than *F_ST_MaxSNP_* (Figure S4). Overall, these analyses provide only modest support for the notion that *F_ST_MaxSNP_* could help detect peaks of genetic differentiation driven by local adaptation that have been narrowed by migration and recombination.

In the above simulations, we used a sample size of 50 chromosomes per population. We generally expect statistical power to be correlated with sample size and understanding the effect of sample size on the power of each statistic is relevant when designing an experiment or choosing which statistics to use. We analyzed the power of *F_ST_MaxSNP_*, *F_ST_Window_*, and *χ_MD_* in three scenarios for high *N_e_* and three for low *N_e_*. We chose scenarios in which *F_ST_MaxSNP_* and the window wide statistics performed differently: a mostly complete soft sweep, a complete soft sweep with a bottleneck, and a partial hard sweep. We found that sample size had a stronger effect on *F_ST_MaxSNP_* than on the window wide statistics (Figure 3). *F_ST_MaxSNP_* is based on allele frequencies at a single site, so it is more sensitive to the increased sampling variance at lower sample sizes than window wide statistics. The sampling variance in each SNP in a window should fluctuate around the mean, so when information from each SNP is combined the window-wide statistic suffers less from the reduced sample size. Demographic history also affected the effect of sample size on each statistic: in scenarios with a population bottleneck, which also increases sampling variance, the power of *F_ST_MaxSNP_* changed from near 1 at sample size 50 or higher to 0 at sample sizes smaller than 50 (Figure 3C, 3D). More generally, *F_ST_MaxSNP_* was found to perform much better with 50 chromosomes than with 20, but showed relatively less improvement for sample sizes larger than 50.

**Figure 3.**
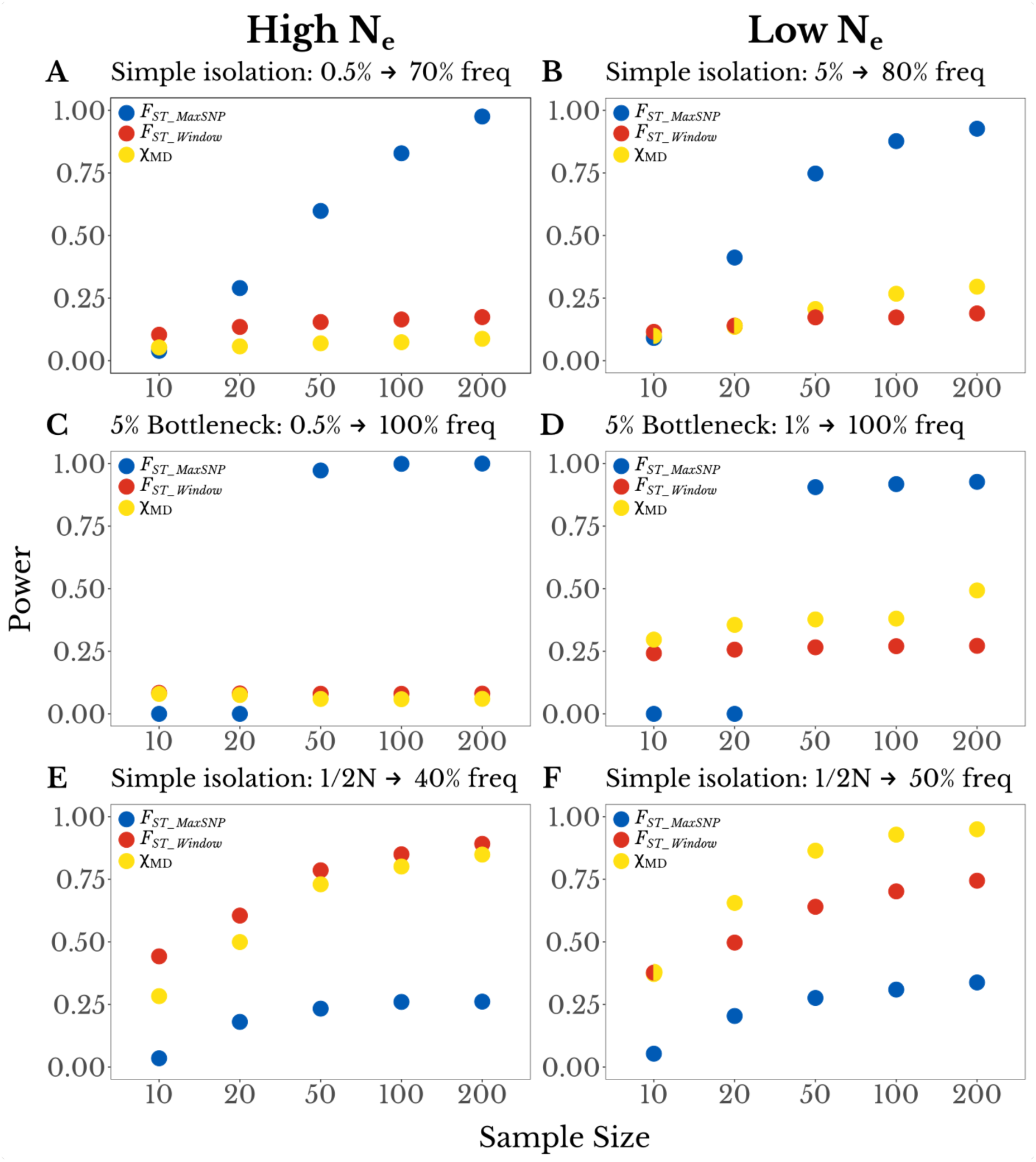
The power of *F_ST_MaxSNP_* is particularly sensitive to sample size. Here, the power of each statistic (y-axis) is plotted as a function of sample size (x-axis; number of chromosomes per population). We found that depending on sample size, *F_ST_MaxSNP_* outperforms *F_ST_Window_* and *χ_MD_* for a simple isolation model, for: (A) a high *N_e_* population with initial beneficial allele frequency of 0.005 and final frequency of 0.70,and (B) a low *N_e_* population with initial frequency 0.05 and final frequency of 0.80. Similar results were observed for a complete soft sweep with a population bottleneck of 0.05, except that the loss of power for *F_ST_MaxSNP_* was more immediate at lower sample sizes, for: (C) a high *N_e_* population with initial frequency 0.05, (D) a low *N_e_* population with initial frequency 0.01. For partial hard sweep scenarios where *F_ST_Window_* and *χ_MD_* outperform *F_ST_MaxSNP_*, all three statistics show more gradual sample size effects, specifically for new mutations and: (E) a final frequency of 0.40 in a high *N_e_* population, and (F) a final frequency of 0.50 in a low *N_e_* population.

We also analyzed the effect of window size on the power of each statistic, with the aim of determining whether there would be a window size for which a single statistic would perform well in contrasting scenarios. For example, one might hope that *F_ST_Window_* for a narrower window might retain good performance for partial hard sweeps, while also capturing the advantages of *F_ST_MaxSNP_* for complete soft sweeps. We explored four scenarios of partial sweeps, two for the high *N_e_* and two for the low *N_e_*. For each population size, we chose one scenario in which the power of *F_ST_MaxSNP_* outperformed *F_ST_Window_* and *χ_MD_*, and one in which it underperformed. In practice, a reduction in window size would result in an increase in the number of tests performed in a genome scan. Therefore, we applied a Bonferroni correction to the p-value proportional to the reduction in size. Our results showed that, for the two scenarios in which *F_ST_MaxSNP_* outperformed *F_ST_Window_* and *χ_MD_*, the power of each statistic remained mostly constant (Figure 4). For the scenarios in which *F_ST_Window_* and *χ_MD_* had an advantage, the power of each statistic, as well as the difference among them, declined with smaller window sizes. Overall, there was no window size in which a single statistic performed well for all scenarios, and hence it may be preferable to apply *F_ST_MaxSNP_* and window-wide statistics separately to empirical data.

**Figure 4.**
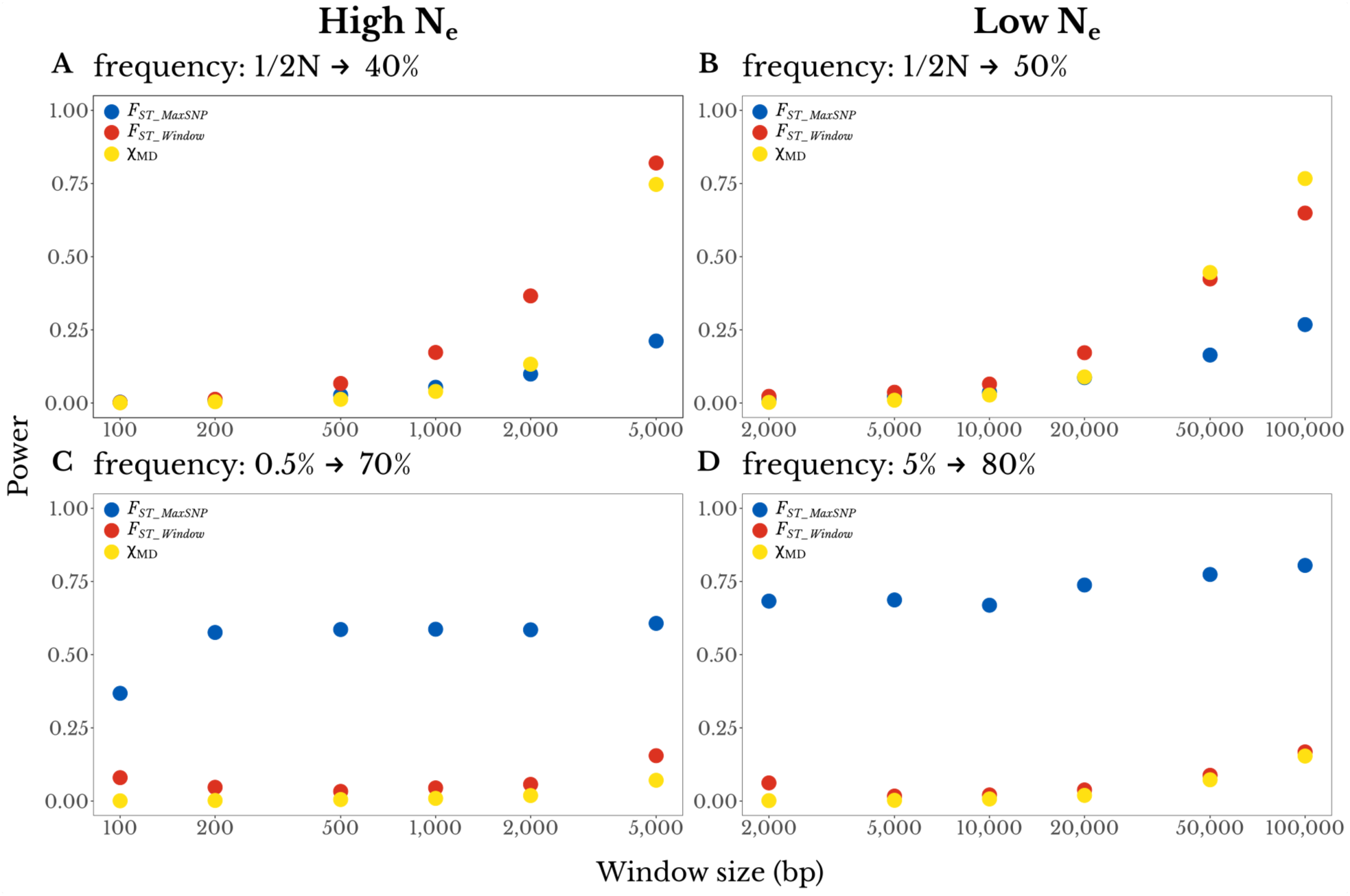
Varying window size does not reveal a single statistic with broad detection power. The top panels show partial hard sweeps for which *F_ST_Window_* and *χ_MD_* outperform *F_ST_MaxSNP_*: (A) a high *N_e_* population with a final beneficial allele frequency of 0.40, And (B) a low *N_e_* population with a final frequency of 0.50. The bottom panels show mostly complete soft sweeps for which *F_ST_MaxSNP_* outperforms *F_ST_Window_* and *χ_MD_*: (C) a high *N_e_* population with an initial beneficial allele frequency of 0.005 and final frequency of 0.70, and (D) a low *N_e_* population with initial frequency 0.05 and final frequency 0.80. These power values reflect a Bonferroni-corrected significance threshold to reflect the relatively larger number of smaller windows needed. Results do not suggest that any statistic in a smaller window size captures the advantages of both *F_ST_MaxSNP_* and the window-wide statistics.

### Outliers for *F_ST_MaxSNP_* and *F_ST_Window_* are enriched in empirical data

In light of the above results, we were interested in applying both *F_ST_MaxSNP_* and *F_ST_Window_* to an empirical data set, in part with an interest in quantifying the relative enrichment of outliers for each statistic, and what that might suggest about the modes of selection active in these populations. We chose to focus on data from the *Drosophila* Genome Nexus (Lack *et al*. 2015, 2016), because it contained several populations of *D. melanogaster* that were linked by an estimated model of population history (Sprengelmeyer *et al*. 2020) and had at least minimal sample sizes available for studying genome-wide patterns of *F_ST_* (Table S2). We included six natural populations of flies. From the ancestral range in Zambia, we included one town population (Siavonga) and one wilderness population (Kafue). We also included four additional town populations: from Rwanda, South Africa, Ethiopia, and France (the latter three having independently colonized colder environments; Pool *et al*. 2017).

We calculated a p-value for each empirical window in each pairwise population comparison, based on neutral distributions of *F_ST_MaxSNP_* or *F_ST_Window_* generated using coalescent simulations of the estimated demographic history (Sprengelmeyer *et al*. 2020; simulation commands in Table S2). Under neutrality, a uniform distribution of p-values is expected. In general, for most population pairs, the distribution of p-values for *F_ST_MaxSNP_* and *F_ST_Window_* showed a U-shape instead of an uniform distribution (e.g. Figure 5A). Nonetheless, the enrichment of high *F_ST_* (defined as p-values from 0 to 0.05) and low *F_ST_* (p-values from 0.95 to 1) varied for each statistic and across the population comparisons (Figure 5B-C).

**Figure 5.**
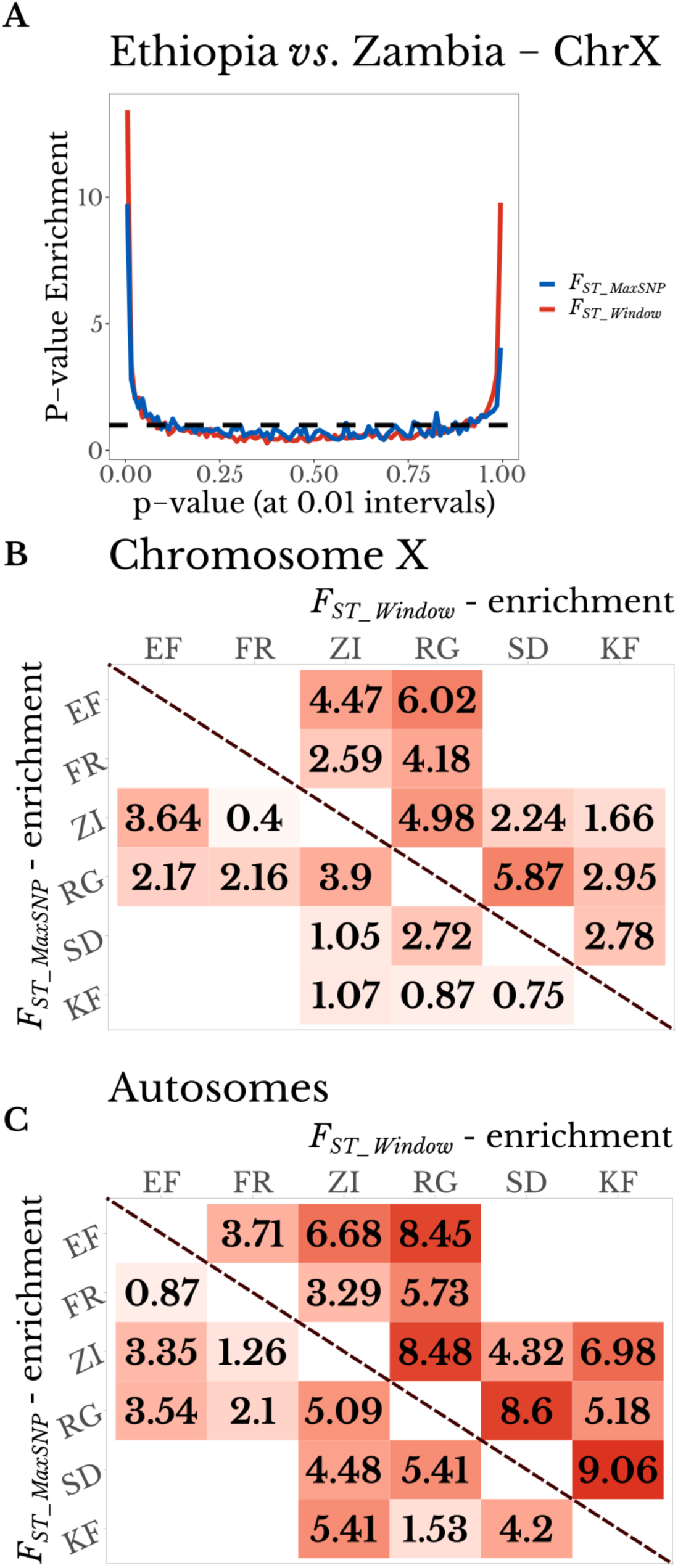
*F_ST_MaxSNP_* and *F_ST_Window_* both show outlier enrichment between natural populations of *D. melanogaster*. (A) Ethiopia-Zambia *F_ST_MaxSNP_* and *F_ST_Window_* values on chromosome X show enrichment of low (right) and especially high values (left), based on the distribution of p-values obtained from neutral demographic simulations. (B and C) *F_ST_MaxSNP_* (lower diagonal) and *F_ST_Window_* (upper diagonal) both show enrichment of high outliers on (B) chromosome X and (C) combined autosome arms. *F_ST_Window_* shows a greater enrichment in nearly all cases. Populations: SD = South Africa. ZI = Zambia. KF = Kafue, Zambia. RG = Rwanda. EF = Ethiopia. Population pairs not present in the same demographic model were not evaluated. Color scale ranges from the minimum to maximum value within each panel.

All population pair comparisons showed an enrichment for windows with high *F_ST_Window_* The smallest enrichment was found between the Zambia (town) and France populations, for which there were 3.29 more windows with high *F_ST_Window_* than expected by chance. The highest enrichment was found in the comparison between the South Africa and Kafue (Zambia wilderness) populations, with an enrichment factor of 9.06. For *F_ST_MaxSNP_*, eight population pairs had an enrichment value > 2, the highest being 5.41 (between the Zambian town and wilderness populations, and between South Africa and Rwanda). On the other hand, one population pair was slightly depleted of windows with high *F_ST_MaxSNP_* (enrichment to 0.87 between France and Ethiopia). In most comparisons, *F_ST_Window_* showed higher enrichment than *F_ST_MaxSNP_*. The only exception was the comparison between South Africa and Zambia (town population), in which both enrichments were very similar: *F_ST_MaxSNP_* enrichment was 4.48 and *F_ST_Window_* enrichment 4.32 (Figure 5). This large variation in enrichment between populations suggests that the kind and prevalence of selective sweeps unique to each population may vary among populations.

The almost universally greater enrichment of *F_ST_Window_* relative to *F_ST_MaxSNP_* could hint that sweeps in the unique detection parameter space of *F_ST_Window_* (*i.e.* partial harder sweeps) are more common among these populations than sweeps in the unique space of *F_ST_MaxSNP_* (*i.e.* more complete softer sweeps). However, the above enrichments may be influenced by locally adaptive sweeps that create multiple linked outlier windows. We therefore pursued a complementary analysis in which nearby outlier windows were merged into “outlier regions”, which were then removed one at a time until the observed enrichment was erased (see Materials and Methods). For almost every population pair, we had to remove a larger number of regions to erase the signal of enrichment of *F_ST_Window_* than the signal of *F_ST_MaxSNP_* (Figure 6). Hence, the greater enrichment of *F_ST_Window_* relative to *F_ST_MaxSNP_* does not appear to be a product of broader linkage signals of *F_ST_Window_* outliers alone.

**Figure 6.**
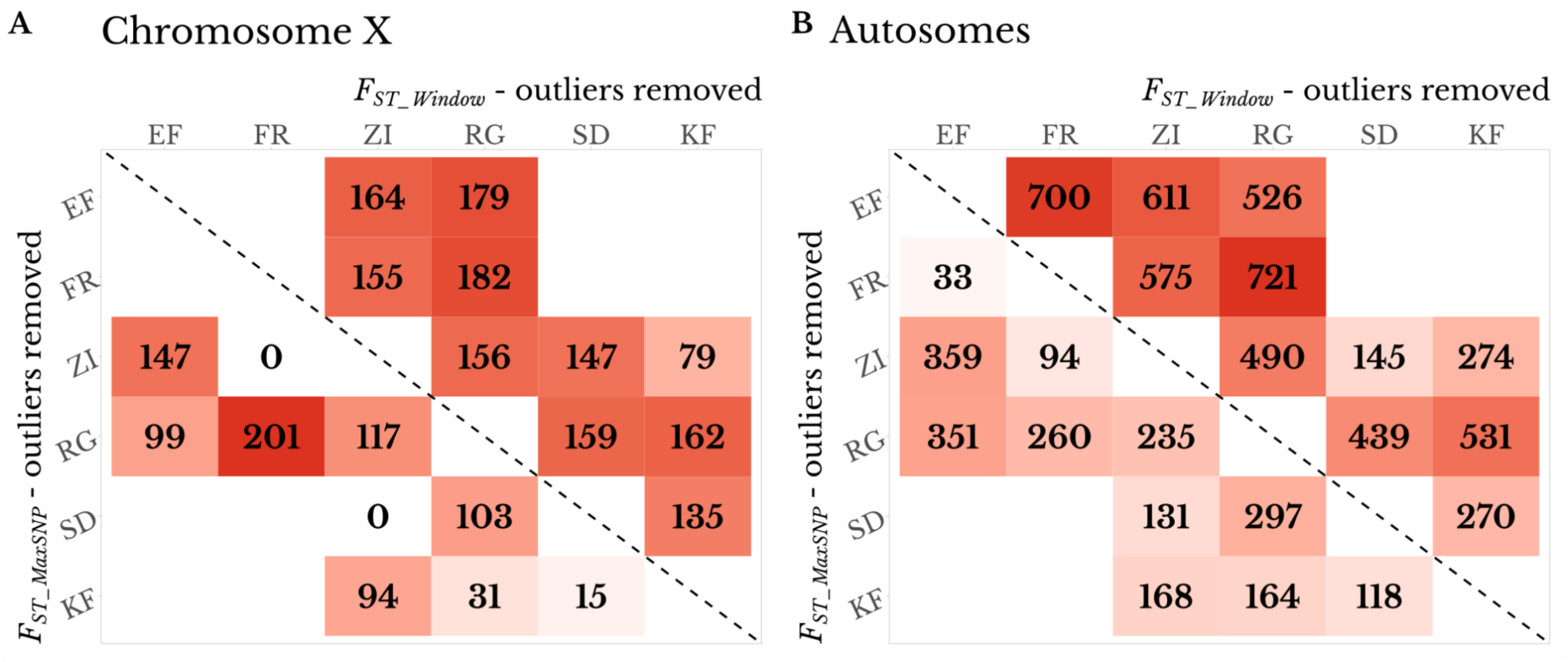
Number of outlier regions that were removed to erase the signature of enrichment for high *F_ST_MaxSNP_* (lower diagonal) and *F_ST_Window_* (upper diagonal) for each population on (A) chromosome X and (B) the combined autosome arms. *F_ST_Window_* was associated with a greater outlier region enrichment for most population pairs, reinforcing the window-level patterns shown in Figure 5. Populations: SD = South Africa. ZI = Zambia. KF = Kafue, Zambia. RG = Rwanda. EF = Ethiopia. Population pairs not present in the same demographic model were not evaluated. Color scale ranges from the minimum to maximum value within each panel.

### Genome Scan for Signatures of Selection

We chose to complement the above multi-population analysis of genome-wide patterns with a closer analysis of a single population pair. We chose to compare the Ethiopia and Zambia town populations because (1) Their relatively large sample sizes of 129-181 and 60-76 respectively for each chromosome arm (Table S2) are more conducive to the analysis of specific *F_ST_MaxSNP_* outliers, (2) These populations showed enrichments of both *F_ST_MaxSNP_* and *F_ST_Window_* (Figure 4), and (3) Past results from these populations helped motivate the present study (*e.g.* Bastide *et al*. 2016). We performed genome scans for regions potentially under population-specific selection between these populations using *F_ST_MaxSNP_*, *F_ST_Window_*, and *χ_MD_*. For each statistic, we obtained a list of outlier windows (top 1%), and as above, we merged nearby outlier windows into regions (Materials and Methods). We obtained 138 outlier regions for *F_ST_MaxSNP_*, 138 for *F_ST_Window_* , and 155 for *χ_MD_*. Our results showed an overlap of just 39% between the outlier regions detected with *F_ST_MaxSNP_* and *F_ST_Window_*. Perhaps surprisingly in light of the above power results, there was a smaller overlap of either *F_ST_* metric with *χ_MD_* (Figure 7A), although the overlap of the haplotype statistic with *F_ST_Window_* was indeed slightly greater. In regions that were outliers for *F_ST_MaxSNP_* but not *F_ST_Window_*, the distribution of individual SNP *F_ST_* values often had a narrow sharp *F_ST_* peak, with most of the other SNPs having low *F_ST_* values. On the contrary, in regions there were outliers for *F_ST_Window_* but not *F_ST_MaxSNP_*, often no single SNP had a large *F_ST_* value, but there was a broad moderate *F_ST_* plateau with many SNPs showing intermediate *F_ST_* values (Figure 8).

**Figure 7.**
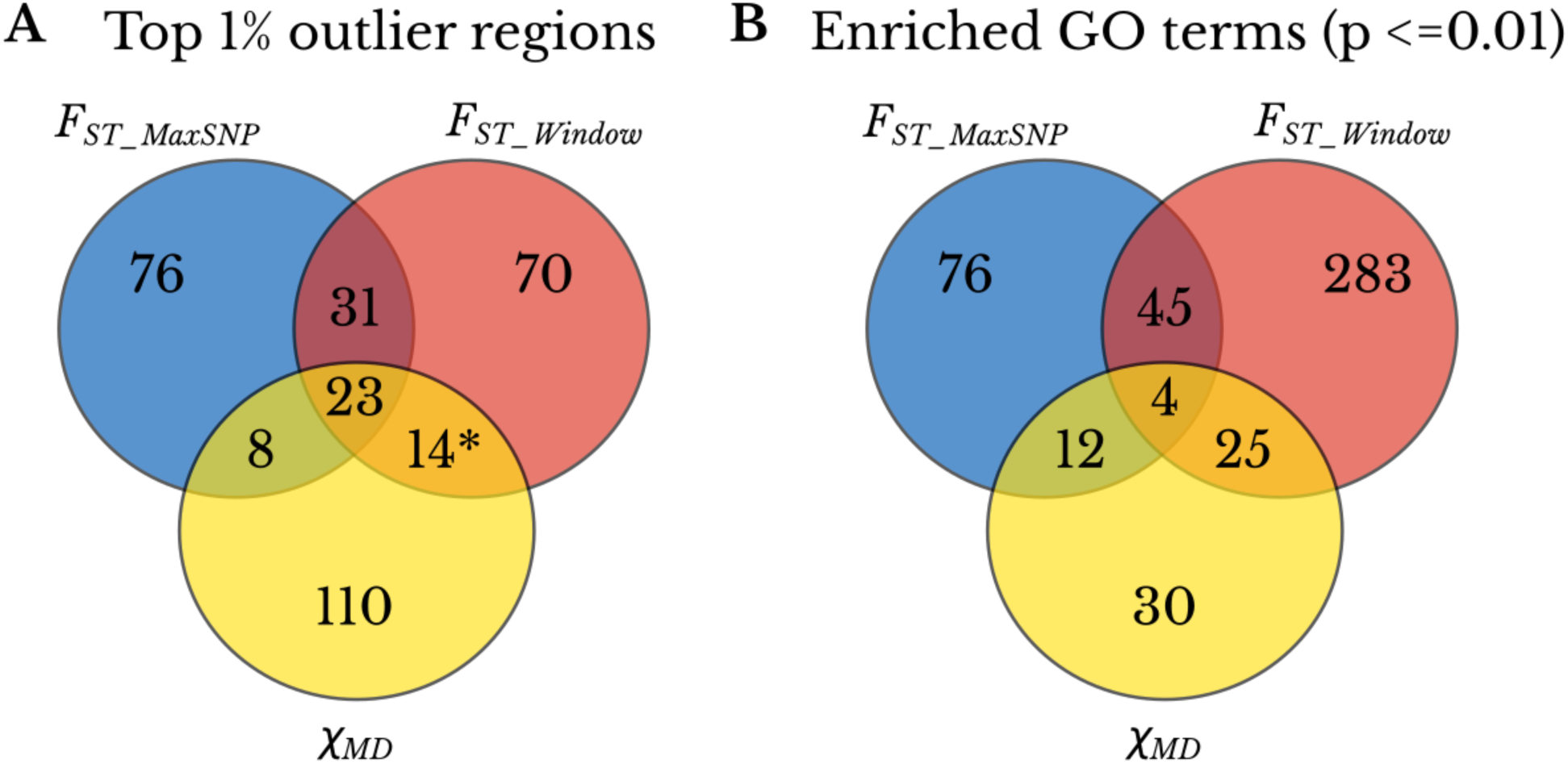
The three statistics detect mostly unique genomic regions and functional categories. (A) Overlap between the top 1% outlier regions detected with *F_ST_MaxSNP_*, *F_ST_Window_*, and *χ_MD_*. * indicates the average number of outlier regions between the two statistics: 15 *F_ST_Window_* outlier regions exclusively overlap *χ_MD_* outliers and 13 *χ_MD_* outlier regions exclusively overlap *F_ST_Window_* outliers. (B) Overlap between enriched GO terms with raw p-value <= 0.01 based on the outlier regions detected with *F_ST_MaxSNP_*, *F_ST_Window_*, and *χ_MD_*.

**Figure 8.**
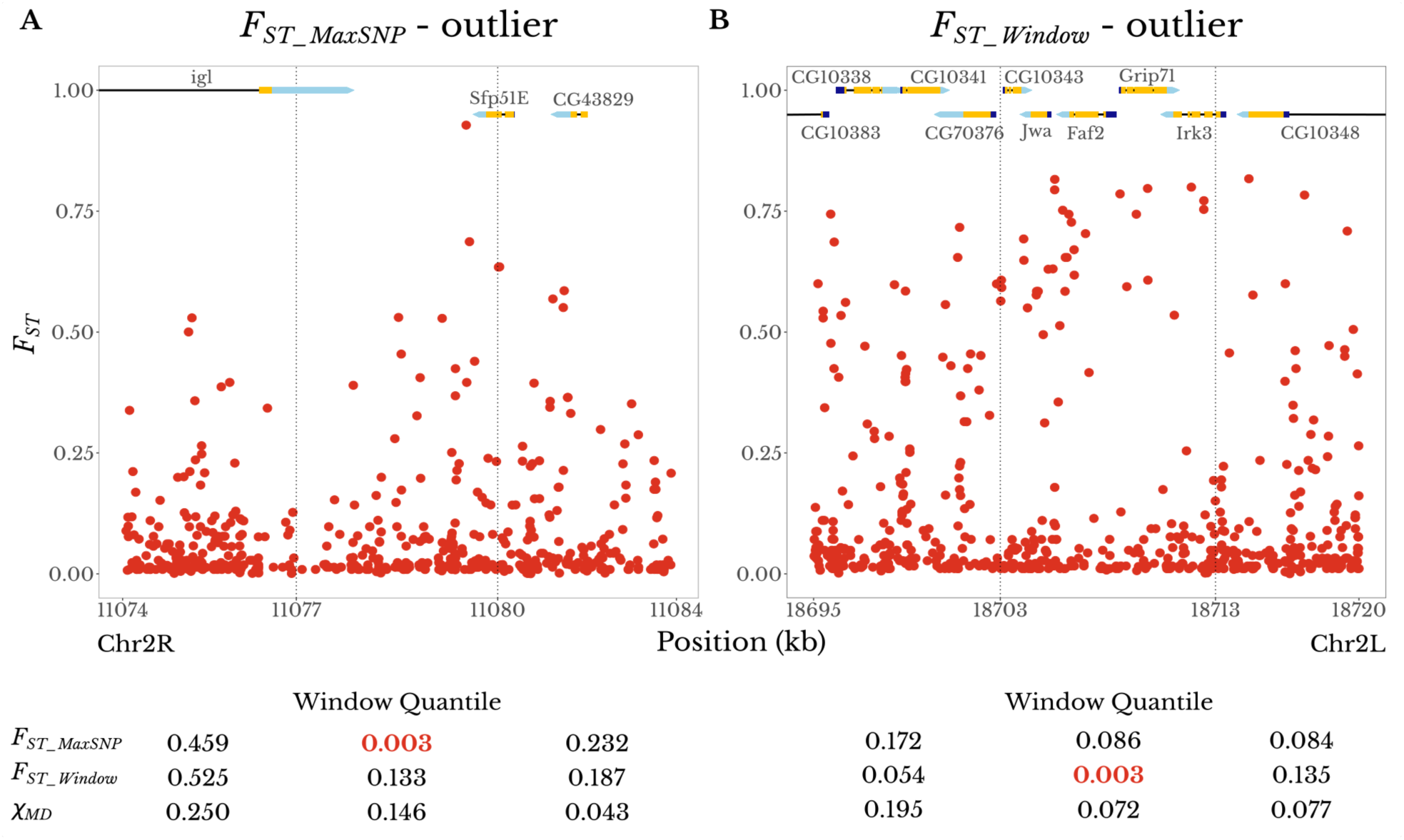
Examples of the distinct SNP-level *F_ST_* landscapes associated with *F_ST_MaxSNP_* versus *F_ST_Window_* outliers. Each plot shows an outlier window for an Ethiopia-Zambia *F_ST_* statistic, plus its adjacent windows. Dashed vertical lines delimit the boundaries of the windows. Numbers under each window are the empirical quantiles of that window’s statistic (*F_ST_MaxSNP_*, *F_ST_Window_*, and *χ_MD_*) in relation to the chromosome arm-wide distribution of the same statistic, with the outlier (quantile < 0.01) value in red. (A) An outlier window for *F_ST_MaxSNP_* (center) shows a peak-like *F_ST_* landscape with one particularly differentiated SNP. (B) An outlier window for *F_ST_Window_* (center) shows a broad plateau of fairly high *F_ST_* values. Gene names and structures are shown at the top of each plot. Protein-coding exons are in yellow, while 5’ and 3’ untranslated regions are in dark blue and light blue, respectively.

We then performed GO term enrichment analysis separately for each statistic’s list of outlier regions. Considering only GO terms with raw p-value < 0.01 from each list, we found mostly lower overlaps between enriched GO terms compared to the spatial overlap between outlier regions (Figure 7B; Table S3). The three statistics differed substantially in the number of enriched GO terms by this criterion: 357 for *F_ST_Window_*, 133 for *F_ST_MaxSNP_*, and 71 for *χ_MD_* (although we emphasize that these terms are not independent and any given list of enriched GO terms will contain overlapping categories). The relative overlap between GO terms enriched for each statistic largely followed the relative numbers of enriched GO terms for each (Figure 7B). Mirroring the outlier region results, most enriched GO terms were detected for only one of the three statistics, consistent with their complementary detection powers described above.

## Discussion

### *F_ST_MaxSNP_* complements other statistics by detecting soft sweeps

Identifying regions under selection can help us answer further questions about the evolution of local adaptation, such as which biological functions are under selective pressure, the number of loci underlying adaptive events, the source of the adaptive variation, and the kinds of genetic changes that might be under selection. Our results underscore the importance of deploying methods capable of capturing different kinds of selective sweeps when the aim of the study is to identify as many genes potentially under local adaptation as possible.

*F_ST_MaxSNP_* in particular, seems to be especially useful to detect soft sweeps with relatively large initial and final frequencies of the beneficial allele. Instances of mostly complete soft sweeps, as simulated here, represent regions in which a beneficial allele was present in several different haplotypes that might have increased in frequency along with the beneficial allele.

While the selected SNP itself changed in frequency drastically, resulting in a large *F_ST_MaxSNP_*, the alleles around it must have changed in frequency to a lesser degree because many background haplotypes were hitchhiking along with the beneficial allele. Therefore, while the beneficial variant can have an extreme *F_ST_* value, the lower allele frequency changes in the other SNPs in that window would result in a *F_ST_Window_* that is not statistically significant, and thus a low power to detect a selective sweep under these conditions.

The window-wide metrics, *F_ST_Window_* and *χ_MD_*, had greater power than *F_ST_MaxSNP_* to detect relatively harder, partial sweeps that had intermediate final allele frequencies. In these sweeps, no individual SNP changed dramatically in frequency, so none have *F_ST_* values higher than what could be obtained randomly in the genome. However, the increase in frequency of one or a few haplotypes resulted in many SNPs in the same region with intermediate *F_ST_*, producing a window-wide pattern that is too extreme to be generated by chance - even if each single marker individually did not have an extreme *F_ST_* value.

There was little difference in the power of *F_ST_MaxSNP_* and *F_ST_Window_* to detect regions under selection in scenarios with varying migration rates. We had expected that *F_ST_MaxSNP_* would outperform *F_ST_Window_* in scenarios with older splits, as selection might only maintain a narrow window of differentiation between the two populations in the presence of long-term recombination with migrant haplotypes. Nonetheless, differences in split time between the two populations only had a small effect in a very narrow space of parameters (intermediate migration rates for high *N_e_* populations, Figure S1), suggesting that even in scenarios with recent divergence, the populations had already reached a state of equilibrium and the balance between migration, selection, and recombination did not result in distinguishable signatures of selection between *F_ST_MaxSNP_* and *F_ST_Window_*. However, both metrics outperformed *χ_MD_* on the simulated scenarios, indicating that selection could not maintain long shared haplotypes in the presence of migration.

In light of the complementary performance of *F_ST_MaxSNP_* and *F_ST_Window_* for the non-migration cases, we tested whether *F_ST_Window_* across shorter windows could yield a balance of reasonable power to detect both complete soft sweeps and partial hard sweeps. However, the relationship between window size and the power - while accounting for the increase in the number of tests in smaller windows - did not follow this prediction. Our results suggest that applying both *F_ST_MaxSNP_* and *F_ST_Window_* to conventionally-sized windows is preferable to shrinking the window size in an effort to identify narrower soft sweeps. More generally, we suggest that genetic differentiation on both SNP and broader scales should be incorporated into scans for local adaptation, whether using the specific summary statistics described here, or attempting to develop a single statistic or integrated analysis framework that encompasses the advantages of both.

In this study, we have used neutral demographic simulations to estimate statistical power at the single window level, only penalizing multiple tests when comparing between window sizes. Clearly, our results do not imply the power to identify genome-wide significant loci, which is only rarely attainable for population genomic scans. Instead, most genome scans aim to identify good candidates for downstream study, and our results are best interpreted in terms of the relative utility of these summary statistics to identify local adaptation candidates.

An important caveat of using *F_ST_MaxSNP_* is that it requires a greater sample size than *F_ST_Window_*. With smaller samples, it is easy to get a large *F_ST_MaxSNP_* at one of the many analyzed SNPs through sampling variance alone, whereas an extreme *F_ST_Window_* value is less likely in this scenario. It is difficult to provide any universal advice regarding sample size, because the neutral variance of *F_ST_MaxSNP_* also depends strongly on demographic history, as shown above. Nonetheless, we have shown that in two scenarios in which *F_ST_MaxSNP_* outperformed *F_ST_Window_* its power declined considerably when we decreased the sample size from 50 to 20 chromosomes. Although the relationship between sample size and power will depend on the specific populations being studied, the utility of *F_ST_MaxSNP_* seems most promising when sample sizes are around 100 alleles per population or more. However, it would be advisable to conduct neutral simulations based on estimated or suspected demography, in order to identify sample sizes for which it is very unlikely to get extreme SNP *F_ST_* values in the absence of local adaptation.

### Both *F_ST_Window_* and *F_ST_MaxSNP_* outliers are enriched among *Drosophila* populations

When we applied *F_ST_Window_* and *F_ST_MaxSNP_* to empirical data from *D. melanogaster* populations, we found that enrichment patterns of *F_ST_Window_* and *F_ST_MaxSNP_* varied among population pairs, both for high and low *F_ST_* values. The excess of windows with high *F_ST_* observed could be explained by local adaptation: unique selective sweeps in one population increase the differentiation between two populations in that region. Not all population pairs showed the same degree of enrichment for high *F_ST_*. A larger enrichment could be due to a higher number of selective sweeps between two populations, stronger selective events that impacted a larger region of the genome, or a neutral history more conducive to outlier detection. The populations we studied cover a large geographical scale, most are located in sub-Saharan Africa and one in Europe. These populations are exposed to a variety of environments, ranging from warm tropical lowlands to cool high latitude and high altitude regions, in addition to commensal versus wilderness settings (Sprengelmeyer *et al*. 2020). Hence, they are most likely exposed to several unique selective pressures that could be underlying local adaptation and an enrichment of high *F_ST_* values.

Alternatively, enrichment for high *F_ST_* could also be explained by background selection, which is expected to reduce genetic diversity and therefore result in lower effective population sizes in that genomic region. Genetic drift is stronger in regions of low *N_e_*, which could increase the differentiation between two populations and produce high *F_ST_* (Charlesworth *et al*. 1993). However, a simulation study of background selection targeting stickleback exons found no evidence for background selection increasing *F_ST_* outliers (Matthey-Doret and Whitlock 2019).

On the other extreme, the existence of enrichment for low values of *F_ST_* suggests that many regions of the genome maintained unexpectedly similar allele frequencies between two populations. Following a population split, neutral evolutionary forces such as genetic drift are expected to increase the genetic differences between two populations. The fact that many regions seemed to have changed less than what was expected due to neutral forces could also be explained by the action of natural selection. This could be the product of shared selective sweeps (i.e. similar selective pressures) taking place in both populations, instead of local adaptation. Shared balancing selection could also be acting at some loci to maintain allele frequencies constant between two populations, perhaps even from before their split time.

We should also acknowledge that the demographic models applied here are simply the best available estimates of population history, and no demographic model fully accounts for the complexity of natural populations. Demographic model misspecification could result in some enrichment of high and/or low *F_ST_* values. One potential source of error in demographic estimation is natural selection. The demographic models were estimated based on tentatively neutral regions of the genome (Sprengelmeyer *et al*. 2020). However, these regions could be under the influence of linked positive and negative selection, with the potential to bias demographic estimation. For example, if the presumed neutral data was substantially affected by either local adaptation or shared sweeps, it could bias the neutral distribution of *F_ST_* towards higher or lower values, respectively, making it more difficult to detect *F_ST_* outliers in that direction. Nonetheless, previous work suggests that this effect might be weak on demographic inference in *D. melanogaster* (Lange and Pool 2018).

In nearly all population pairs, *F_ST_Window_* showed a larger enrichment than *F_ST_MaxSNP_*. The greater enrichment of *F_ST_Window_* persisted when we instead pursued an outlier region removal strategy. In light of the complementary zones of power shown in Figure 1, these results suggest that roughly speaking,there might be a larger contribution of partial hard sweeps than complete soft sweeps to local adaptation among these populations. Furthermore, the fairly low levels of outlier overlap between *F_ST_Window_* and *F_ST_MaxSNP_* may suggest that the sweeps both statistics can reliably detect (*i.e.* more complete harder sweeps) are not the primary drivers of local adaptation in this data set. Overall, these results suggest that partial sweeps might have played a large role in the adaptation of fly populations to diverse environments. The importance of partial sweeps in populations of *D. melanogaster* has been proposed previously, including for some of the populations studied here (Pool and Aquadro 2007; Bastide *et al*. 2016; Garud and Petrov 2016; Vy *et al*. 2017).

Here, we have shown that SNP-level *F_ST_* (*F_ST_MaxSNP_*) offers strong power to detect soft sweeps, and is highly complementary to window-wide frequency and haplotype statistics for detecting local adaptation. These results stress the importance of taking into account the different signatures left by different kinds of selective sweeps in the genome when deciding how to perform a genome scan. The raw summary statistics evaluated here can either be applied in parallel, or their signals can be integrated into frameworks such as approximate Bayesian computation and machine learning. Thus far, the latter methodologies have been used more extensively to detect and classify selective sweeps within a single population (Peter *et al*. 2012; Sheehan and Song 2016; Schrider and Kern 2016, 2017). However, such approaches are equally applicable to the study of local adaptation (Key *et al*. 2014). Future work could investigate whether methods that combine multiple statistics would benefit from including *F_ST_MaxSNP_*, potentially increasing their power to detect soft sweeps and their accuracy in classifying different types of sweeps. Because studies of genetic differentiation between populations inherently control for evolutionary variance in the shared ancestral population, local adaptation may offer a better “signal to noise ratio” regarding the types of positive selection acting in natural populations, compared to single population studies. Hence, our results may contribute toward not only an improved ability to detect local adaptation, but also a clearer understanding of adaptation in nature more generally.

## Methods

### Simulation Power Analysis

To generate adaptive and neutral distributions of genetic diversity, we performed simulations of demographic history scenarios with and without natural selection using *msms* (Ewing and Hermisson 2010). For each model, we obtained 10,000 replicates from which we calculated the statistics of interest. Power was calculated as the proportion of replicates under selection with a statistical value larger than 95% of the values obtained in its corresponding replicates without selection. We investigated the power of three different statistics: *F_ST_MaxSNP_*, *F_ST_Window_* and *χ_MD_* (Lange and Pool 2016), which were calculated on windows of fixed size. *F_ST_MaxSNP_* is based on the SNP within a window with the highest *F_ST_* value. *F_ST_Window_* was calculated as the weighted average of all SNPs in a window (Reynolds *et al*. 1983). *χ_MD_* stands for Comparative Haplotype Identity; it compares the average length of identical haplotypes in a window between two populations, and was calculated following Lange and Pool (2016). The window size used was 5,000 bp for simulations of populations with high effective population size (*N_e_*) and 100,000 bp for simulations of populations with low *N_e_*. Except where otherwise stated, the sample size was 50 chromosomes. The high *N_e_* simulations used parameters similar to those from flies (*Drosophila melanogaster*) while the low *N_e_* had parameters similar to humans (simulation parameters followed Lange and Pool, 2016).

We initially used scenarios of constant population size and a simple population split to simulate scenarios of selective sweeps with varying initial and final allele frequencies, representing hard and soft sweeps as well as complete and partial sweeps. We also simulated scenarios of population bottlenecks and population splits for complete selective sweeps, and for scenarios with varying migration rates for hard sweeps (not constrained by ending allele frequency). For bottlenecks, the population that will experience local adaptation underwent a period of reduced population size for the first 0.01 coalescent units after the population split (which in most scenarios including these, occurred 0.05 coalescent units ago; Table S1).

The simulations of populations with high *N_e_* were done for two different selection coefficients (s = 0.01 and s = 0.001) and simulations of populations with low *N_e_* only included s = (Table S1). Simulations of complete sweeps only used replicates in which the beneficial allele went to fixation. Simulations of partial sweeps only accepted replicates in which the beneficial allele stayed within 4% of the targeted ending frequency. Selection initiation time was adjusted in each case to maximize the proportion of accepted replicates. Moreover, in the scenarios with initial allele frequencies larger than 1/2*N_e_*, both the selected and non-selected populations had the same initial frequency.

For models that included migration (gene flow), selection of equal magnitudes but in opposite directions was imposed on each population. Per generation migration rates varied from 0.0004 to 0.004 in simulations with high *N_e_* populations and from 0.01 to 0.10 in simulations with low *N_e_* populations. For each migration rate, split times varied from 0.1 to 1 coalescent unit.

We calculated the effect of sample size on the power of each statistic in six different scenarios: four models with demographic history of a simple isolation between two populations and two models with population size bottleneck. Of the simple isolation models, two models for high *N_e_* populations were considered: one in which *F_ST_Window_* outperformed *F_ST_MaxSNP_* (initial allele frequency of 1/2*N_e_* and final allele frequency of 0.4) and another where *F_ST_MaxSNP_* outperformed *F_ST_Window_* (initial frequency of 0.005 and final frequency of 0.7). Two scenarios for low *N_e_* populations were also considered: one in which *F_ST_Window_* outperformed *F_ST_MaxSNP_* (initial allele frequency of 1/2*N_e_* and final allele frequency of 0.5) and another where *F_ST_MaxSNP_* outperformed *F_ST_Window_* (initial frequency of 0.05 and final frequency of 0.8). For the bottleneck models, we used models with a bottleneck of 5% (*i.e.* a reduction to 5% of the prior *N_e_* for 0.01 coalescent units in the adapting population immediately following the population split) and only models in which *F_ST_MaxSNP_* outperformed the window wide statistics were considered: one model for high *N_e_* population (initial allele frequency from 0.5% to 100%) and one for low *N_e_* populations (initial allele frequency from 1% to 100%). For all the six scenarios, we used sample sizes of 10, 20, 50 (original sample size), 100, and 200 chromosomes.

We calculated the effect of window sizes on the power of each statistic in four different scenarios, the same scenarios of simple isolation used to calculate the power of sample sizes above. For the high *N_e_* scenarios, we used window sizes of 5 kb (original size), 2 kb, 1 kb, 0.5 kb, kb, and 0.1 kb. For the low *N_e_* scenarios, we used window sizes of 100 kb (original size), 50 kb, 20 kb, 10 kb, 5 kb, and 1 kb. To calculate *χ_MD_*, we used a minimum haplotype threshold of 10% of the window size (as was used for the original analyses). For each window size smaller than the original, we applied a p-value Bonferroni multiple testing correction proportional to the reduction in size (or equivalently, the increased number of windows needed to cover a given genomic region) to calculate power. That is, while for the standard window size power is the number of replicates with a p-value of 0.05 or lower, for a window half the size of the original the p-value would need to be 0.025 or lower.

### Empirical Enrichment of *F_ST_MaxSNP_* and *F_ST_Window_* - data and simulations

Our data set consists of individual fly strain genomes from six natural populations of *D. melanogaster*: one non-human commensal population from Kafue, Zambia (KF) and five human commensal populations from different countries: Zambia (ZI), South Africa (SD), Rwanda (RG), Ethiopia (EF) and France (FR), using data from Lack *et al*. (2016) and Sprengelmeyer *et al*. (2020). From each population, for each chromosome arm (ChrX, Chr2L, Chr2R, Chr3L, Chr3R), we excluded genomes from lines with a known inversion for that arm. To boost the sample size of two populations with genomes from partially inbred lines (Ethiopia and France), instead of only using homozygous regions of the genome (as in the original filtering of the published data set) we also included heterozygous regions identified by Lack *et al*. (2016), and therefore counted two alleles at each site from these regions. For any pair of lines with excess identity by descent (IBD) between them (defined as more than 10 megabases of IBD outside previously defined regions of low recombination; Lack *et al*., 2016), we excluded one member of the pair from this data set. For each population sample and each chromosome arm, we chose a sample size to jointly maximize the number of analyzable sites and the sample size itself. Our resulting sample sizes are shown on Table S2. For sites with more than that number of alleles called, we downsampled to match the chosen sample size.

We calculated pairwise *F_ST_Window_* and *F_ST_MaxSNP_* for all populations using diversity-scaled window sizes designed to contain 250 non-singleton SNPs in the ZI sample. To compare empirical and null distributions for similar recombination rates, each window was assigned to one of five recombination rates bins based on estimates from Comeron *et al*. (2012); the bins corresponded to recombination rates from 0.5-1, 1-1.5, 1.5-2, 2-3, and greater than 3. Windows with recombination rates lower than 0.5 were not used due to low spatial resolution for localizing signatures of selection in low recombination regions. We obtained p-values for each window using neutral demographic simulations performed using *ms* (Hudson 2002).

Demographic simulations were performed using parameters estimated for the evolutionary history of nine populations of *D. melanogaster*, including all the populations we analyzed (Sprengelmeyer *et al*. 2020). The other three populations were lowland Ethiopia (EA), Cameroon (CO), and Egypt (EG). We did not use those three populations in our empirical analyses due to their lower sample sizes. Nonetheless, they were included in the simulations in order to accurately reflect the estimated patterns of migration.

Each demographic model had been estimated based on tentatively neutral genetic markers (short introns and 4-fold synonymous sites from regions with sex-averaged recombination rates of at least 1 cM/Mb) from inversion-free chromosome arms (Sprengelmeyer *et al*. 2020). A model was estimated for each of three chromosome arms that had lower inversion frequencies (X, 2R, and 3L), and the history was inferred iteratively, such that not all population samples were present in the same model. To better approximate genetic diversity in all populations, we used two sets of demographic models: Northern model (containing ZI, RG, CO, EF, FR, EG, EA) and Southern model (containing ZI, RG, CO, SD, and KF).

The Northern model for the chromosome X was subdivided into two sub-models (one with ZI, RG, CO, EF, EA and another with ZI, RG, CO, FR, EG). Hence, we simulated four Northern models and three Southern models (command lines in Table S2). The models for the autosomal chromosome arms (2R and 3L) were simulated using the highest sample sizes for any autosomal arm of each population (Table S2). Simulated sample sizes were downsampled to match the sample sizes of each specific arm when comparing empirical and simulated *F_ST_* patterns for any given arm. The window size and crossing over rate used in each replicate were based on a random sampling with replacement from the empirical windows, and the single gene conversion rate and mean tract length were based on the estimates of Comeron *et al*. (2012). Therefore, a null distribution was generated for each model and each recombination bin (described above). For each model and each recombination bin, 50,000 replicates were simulated.

### Enrichment calculation

*F_ST_Window_* and *F_ST_MaxSNP_* were calculated for each population pair and each chromosome arm. *F_ST_* was calculated for the simulated data using the same sample sizes as the empirical data (Table S2). For sites with more than two alleles, only the two most common alleles were kept. Sites with minor allele counts lower than two were discarded from empirical and simulated analyses.

P-values were calculated for each window based on the neutral distribution of its corresponding recombination group. Windows from chromosome X were compared to neutral distributions based on the model for chromosome X. For autosomal loci, we determined that simulations from the 3L model yielded somewhat milder outlier enrichments than the 2R model, and therefore we conservatively focused on results from the 3L model.

We calculated p-value enrichments for *F_ST_Window_* and *F_ST_MaxSNP_* using p-value bins of width equal to 0.05, resulting in 20 bins of p-value 0 to 1. We counted how many windows had a given p-value for each bin and divided the observed number by how many windows we expected to have with a p-value in that bin based on simulated data.

Neighboring windows with low p-value could be showing the effect of a single selective sweep. Therefore, we complemented this outlier window enrichment analysis with one based on “outlier regions”. We intentionally defined outlier regions generously, preferring to falsely lump two sweeps versus splitting a single sweep into two or more regions. Formally, starting with the window containing the lowest p-values, we extended the region surrounding it until we reached a stretch of five consecutive windows with p > 0.1 to create an outlier region. We removed the outlier regions from our analysis and repeated the process until the signal of enrichment was erased (defined as the p < 0.05 bin having no more enrichment than the 0.05 < p < 0.1 bin). For each of *F_ST_MaxSNP_* and *F_ST_Window_*, we recorded the total number of outlier regions that had to be removed for a given population pair.

### Genome scan for regions under selection - Ethiopia vs. Zambia

We performed a genome scan for candidate regions under selection between the Ethiopia (EF) and Zambia (ZI) populations. We calculated *F_ST_Window_*, *F_ST_MaxSNP_*, and *χ_MD_* for each window of the genome. We used an outlier approach and considered windows in the top 1% of each statistic to be the candidate regions under selection. Here, we combined multiple outlier windows into the same outlier region if they were separated by no more than five windows with p-value > 0.01. To investigate whether the candidate regions detected with each statistic were the same or unique, we calculated how many regions overlapped between the different statistics. We considered that two regions were overlapping if at least 50% of the smaller region overlapped the larger one.

For each list of candidate regions under selection, we performed a GO term enrichment analysis using a method initially described by Pool *et al*. 2012. For each gene within a candidate region, we obtained GO term annotations from FlyBase. The GO terms for each gene also included all the parents of each term. GO terms that appeared repeatedly in a candidate region were counted only once for that region. We calculated the p-values for each GO term based on 10,000 permutations of the genomic locations of the outlier regions. This procedure allows genes to have different null probabilities of being outliers, particularly based on their length. We obtained a list of enriched GO terms for each statistic defined as the GO terms with raw p-values less than or equal or to 0.01. We then determined the overlap between the three lists of enriched GO terms.

### Data Availability Statement

No new empirical data were generated for this research. Scripts used in the analyses presented can be found at https://github.com/ribeirots/fst_maxsnp.git.

## Acknowledgments

We appreciate comments from multiple Pool lab members on this manuscript. This research was funded by NIH grants R01 GM127480 and R35 GM13630, and by NSF grant DEB 1754745.

