## Supplemental Figures for "SNP-level *F_ST_* outperforms window statistics for detecting soft sweeps in local adaptation"

### Supplementary Figures

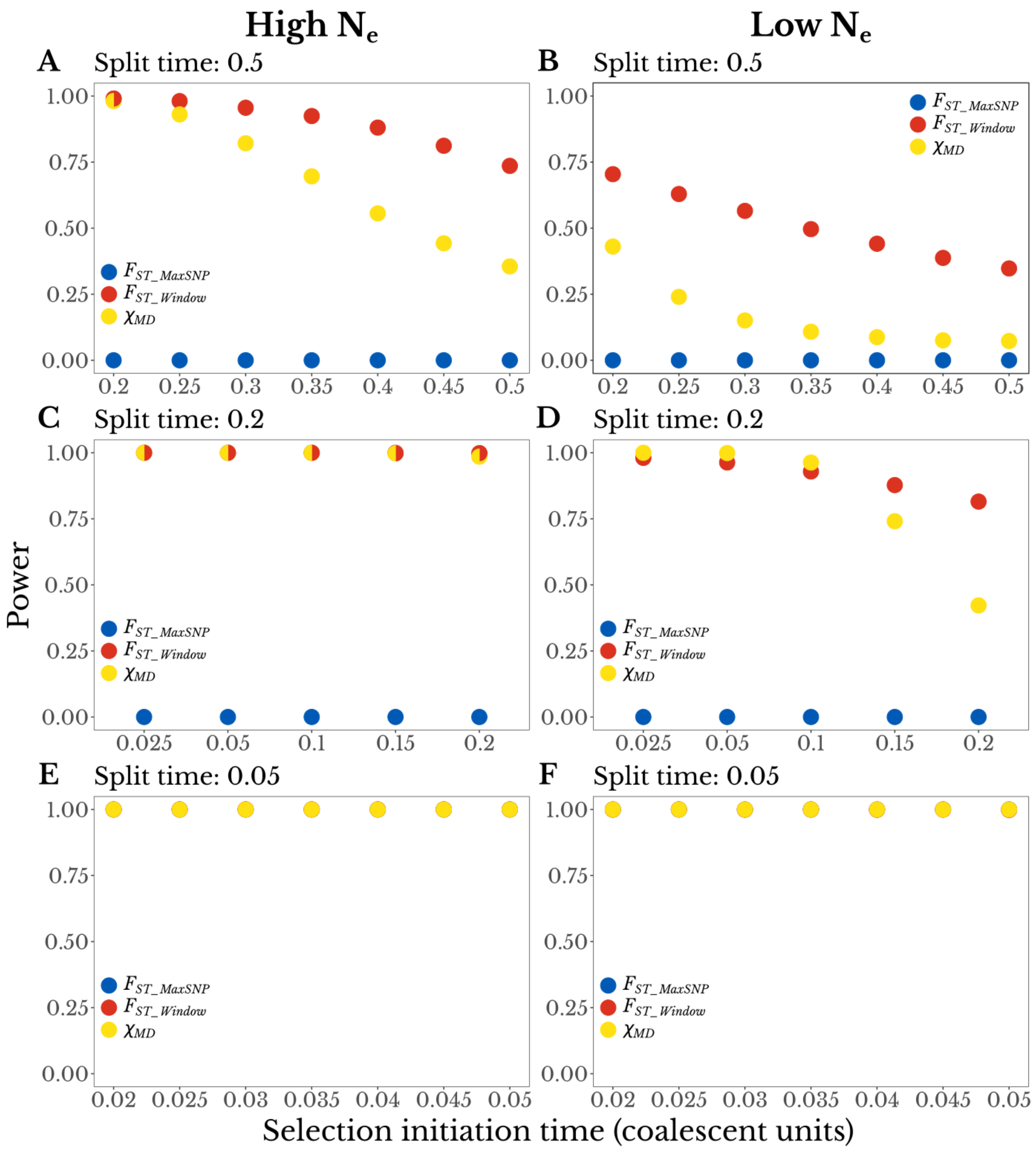


**Figure S1.** Power to detect local adaptation from simulations of complete hard sweeps with varying population split and selection initiation times. Here, the initial frequency of the favored variant is $1/2N_e$

and the final frequency is 1.0. The power of $F_{ST\_MaxSNP}$ was found to be binary, either 0 or 1, depending on whether a given history generated fixed differences in more than 5% of window replicates under neutrality. The power of $F_{ST\_Window}$ and $\chi_{MD}$ also declines with increasing split time (albeit more continuously), and also with increasing selection initiation time. Split time and selection initiation time are measured in coalescent time units (of $4N_e$ generations). These simulations were performed under a simple isolation model between two populations without population size changes or migration. The left column shows results for simulations with high $N_e$ (s=0.001) and the right column shows simulations with low $N_e$ (s=0.01). (A and B) simulations with split time = 0.5. (C and D) simulations with split time = 0.2. (E and F) simulations with split time = 0.05.

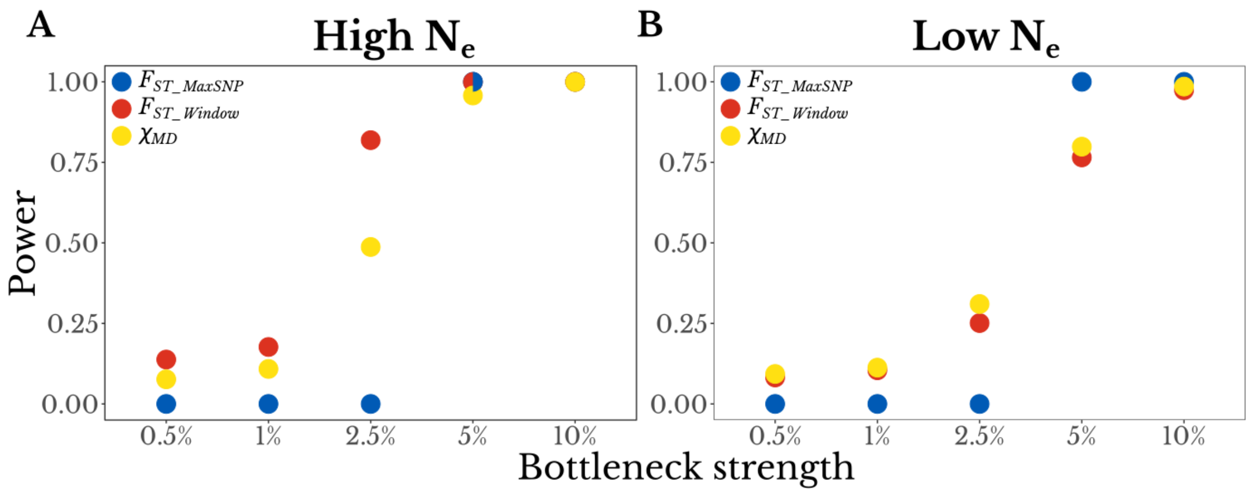


**Figure S2.** Power to detect local adaptation from simulations of complete hard sweeps with a population bottleneck in the adapting population. Bottleneck strength is defined as the proportional redution in population size for the adapting population for the first 0.01 coalescent units after the population split. Similar to Figure S1, the power of $F_{ST\_MaxSNP}$ is binary, either 0 or 1, depending on whether the bottleneck is strong enough to generate fixed differences in more than 5% of window replicates. The power of $F_{ST\_Window}$ and $\chi_{MD}$ is also dependent on bottleneck strength, but it changes gradually. Here, the split time was 0.05 coalescent units and the selection initiation time was 0.025 coalescent units. (A) High $N_e$ populations with s=0.001. (B) Low $N_e$ populations with s=0.01.

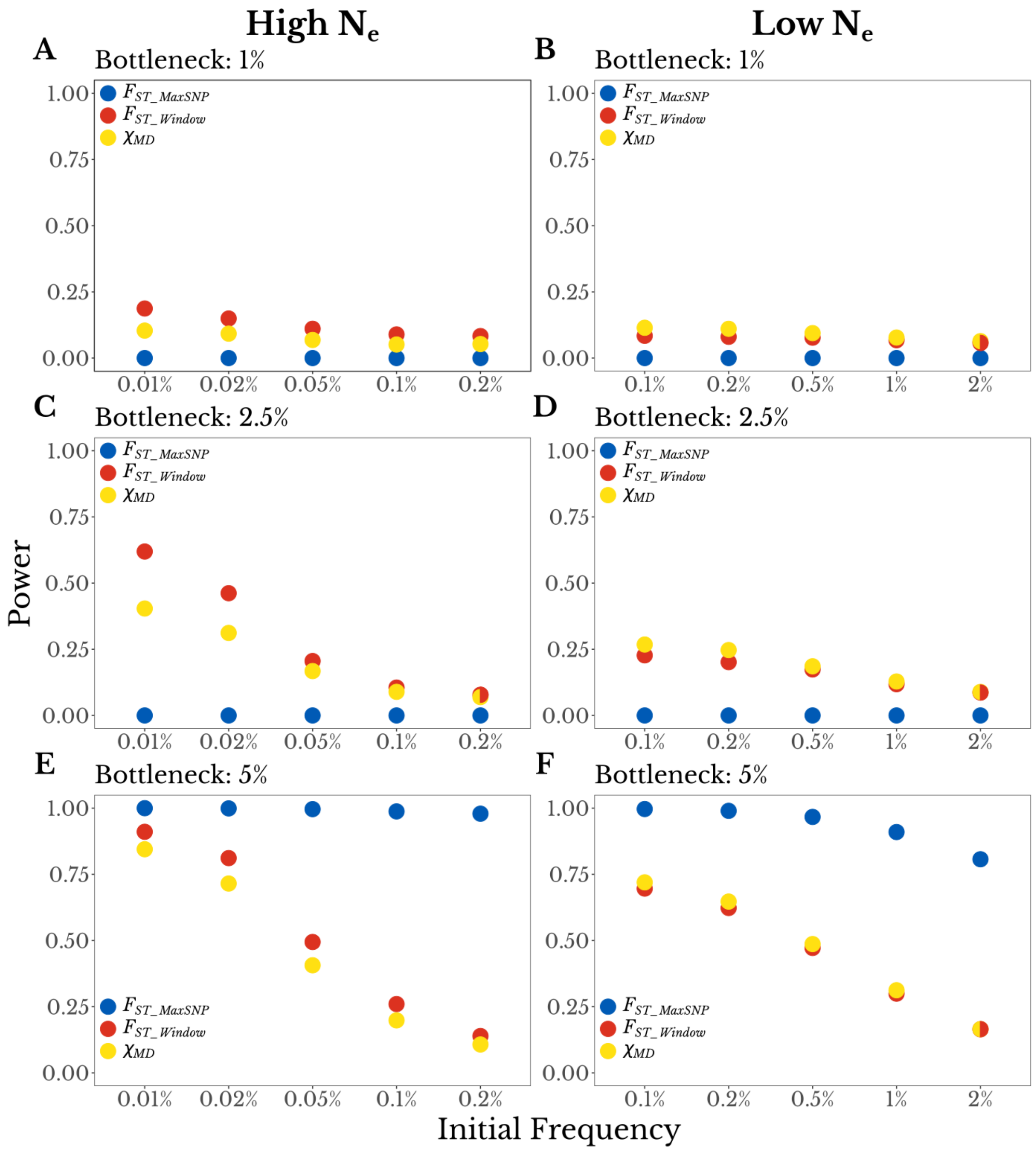


**Figure S3.** Power to detect local adaptation from simulations of complete soft sweeps with a population bottleneck in the adapting population. As in Figure S2, bottleneck strength is defined as the proportional redution in population size for the adapting population for the first 0.01 coalescent units after the population split. The x-axis shows the pre-selection frequency of the favored variant, where larger frequencies are expected to produce softer sweeps that maintain more unique haplotypes. Results indicate that $F_{ST\_MaxSNP}$ outperforms $F_{ST\_Window}$ and $\chi_{MD}$ under weaker population size bottlenecks (see 5% above) and was less affected by higher initial frequencies. However, the power of $F_{ST\_MaxSNP}$ was 0 under stronger bottlenecks that produce fixed differences under neutrality at non-trivial rates. Contrary to Figure S1 and Figure S2, the power of $F_{ST\_MaxSNP}$ is not binary (0 or 1) because both populations start with the same initial allele frequencies, and hence the favored variant is not always observed as a fixed difference. The power of $F_{ST\_Window}$ and $\chi_{MD}$ also increase with weaker bottlenecks, showing more continuous change. The window-wide statistics were relatively more sensitive to higher initial frequencies, consistent with the results shown in Figure 1 and Figure 2 for non-bottleneck cases. As in Figure S2, the split time was 0.05 coalescent units and the selection initiation time was 0.025 coalescent units.

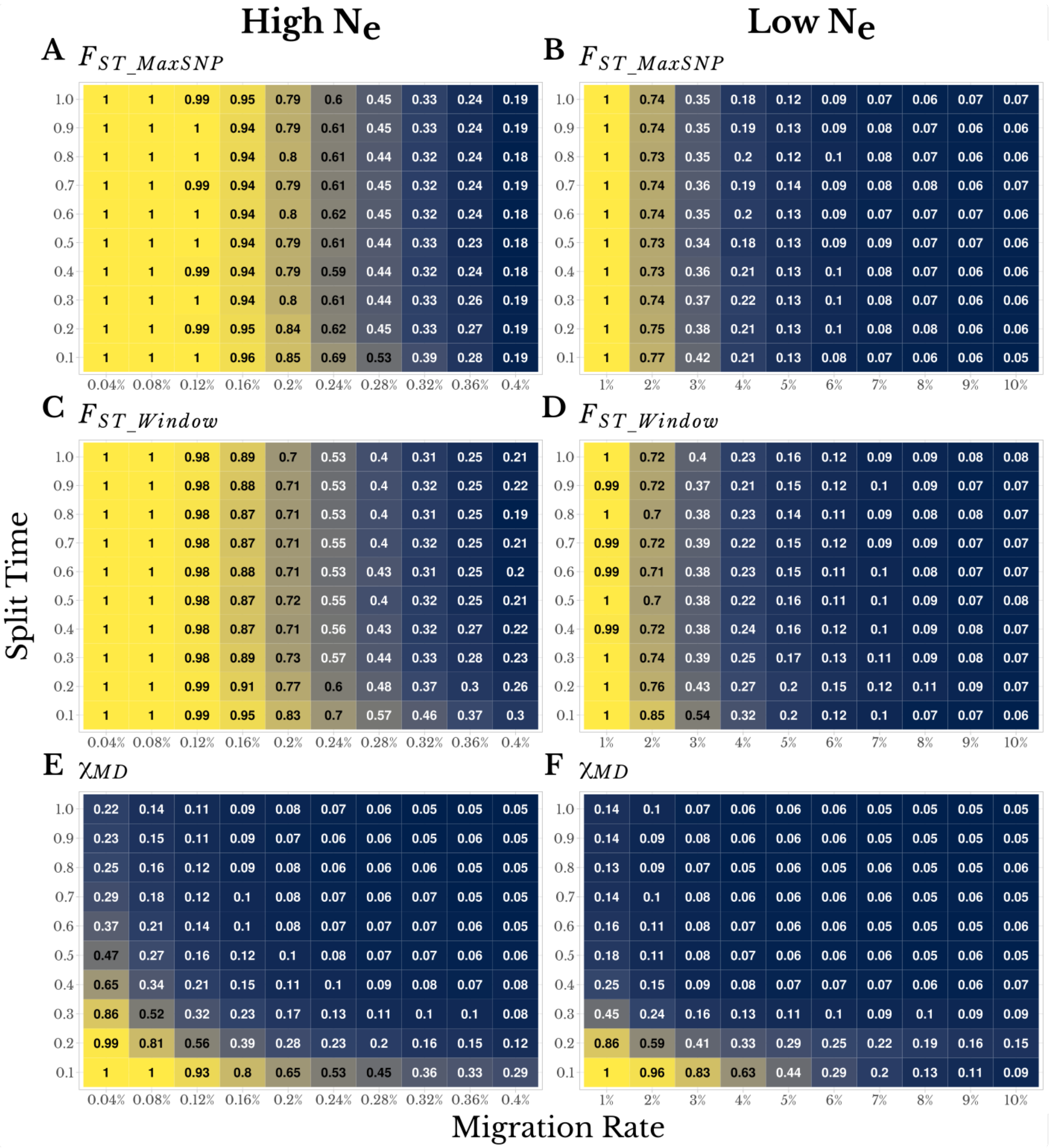


**Figure S4.** $F_{ST\_MaxSNP}$ and $F_{ST\_Window}$ perform similarly in detecting local adaptation from isolation-migration histories, whereas $\chi_{MD}$ shows weaker performance. Numbers and colors in each panel both depict statistical power to detect local adaptation, in high $N_e$ populations (s=0.001, left column) and low $N_e$ populations (s=0.01, right column).  In each panel, the x-axis illustrates the per-generation migration probability in each direction and the y-axis illustrates the population split time in coalescent time units.  These simulations reflect selection on new mutations (initial frequency ($1/2N_e$) and the final frequency varied depending on selection and migration. High $N_e$ populations with s=0.001 are shown in the left column, while low $N_e$ populations with s=0.01 are in the right column.  Power to detect local adaptation is shown for (A and D) $F_{ST\_MaxSNP}$, (B and E) $F_{ST\_Window}$, and (C and F) $\chi_{MD}$.
